# Novel genome assemblies and Evolutionary Dynamics of North American *Anopheles* mosquitoes

**DOI:** 10.1101/2021.08.31.458407

**Authors:** Cory A. Henderson, Karen Kemirembe, Sage McKeand, Christina Bergey, Jason L. Rasgon

**Affiliations:** Departments of Entomology and Disease Epidemiology, The Pennsylvania State University, University Park, PA, United States of America, 16801; Department of Genetics, Rutgers University, New Brunswick, NJ, United States of America, 08901

**Keywords:** Anopheles, Evolutionary Genomics, North American Vectors, Novel Genome Sequences

## Abstract

*Anopheles* mosquitoes are the principal vectors for malaria and lymphatic filariasis, and evidence for arboviral transmission under laboratory and natural contexts has been demonstrated. Vector management approaches require an understanding of the ecological, epidemiological, and biological context of the species in question, and increased interest in gene drive systems for vector control applications has resulted in an increased need for genome assemblies from understudied mosquito species. While the genomes for many *Anopheles* species have been sequenced, North American *Anopheles* taxa have been neglected. In this study we present novel genome assemblies for the North American species *An. crucians, An. freeborni, An. albimanus*, and *An. quadrimaculatus*, and examine the evolutionary relationship between these species. We identified 790 shared single copy orthologs between the newly sequenced genomes and created a phylogeny using 673 of the orthologs, identifying 525 orthologs with evidence for positive selection on at least one branch of the phylogeny. Gene ontology terms such as calcium ion signaling, histone binding, and protein acetylation were identified as being biased in the set of selected genes. These novel genome sequences will be useful in developing our understanding of the diverse biological traits that drive vectorial capacity in anophelines.

## INTRODUCTION

Vectors are living organisms which can transmit disease causing agents between vertebrate hosts. Mosquitoes transmit disease-causing pathogens during the act of bloodfeeding, which is required by many species for reproduction [1]. Vector-borne pathogens generally fall into the broad categories of parasites, bacteria, or viruses, and result in over 700,000 deaths per year, primarily in tropical and sub-tropical regions and disproportionately affecting the poorest populations residing in these areas [2].

*Anopheles* mosquitoes are the principal vectors for malaria and lymphatic filariasis, which historically impacted both North and South America, but now primarily circulates in central and South America [3]. *Anopheles* mosquitoes do demonstrate limited evidence for viral transmission; in natural contexts *Anopheles gambiae* and An. *funestus* act as the primary transmitting vectors for O’nyong’nyong virus, and in laboratory contexts multiple *Anopheles* species have demonstrated capacity for transmission of Mayaro and Chikungunya virus [4-6].

Vector management approaches require an understanding of the ecological, epidemiological, and biological context of the vector species, and with the increase in interest and application of genetic strategies such as gene drives utilizing CRISPR-Cas9, the availability of genomic sequences of vector species is becoming an immensely important aspect of vector control. High quality chromosome level genome assemblies exist for certain model anopheline species such as *An. gambiae* and *An. stephensi* [7-9], however, most *Anopheles* species complexes are understudied and as such comprehensive comparative genomics studies on these species are difficult to perform. Currently, while native anopheline species ranges span the entirety of the continental United States, genome assemblies for of these species are absent from the literature. The primary anopheline species in the U.S. are *An. freeborni* in the west and *An. quadrimaculatus* in the east, and members of the *An. punctipennis* species group such as *An. crucians* are also present throughout the eastern half of the country [3]. *An. albimanus* is one of the primary anopheline mosquitoes in Mexico, Panama and the Caribbean [3], but does occur locally Florida as well [10].

In this study we present novel genome assemblies for the North American anopheline species *An. crucians, An. freeborni, An. quadrimaculatus*, and *An. albimanus*. We examine the evolutionary relationship between these species and identify examples of selection occurring in shared single copy orthologs identified within these novel genome assemblies. Of 790 total identified single copy orthologs shared between all the newly sequenced genomes, 525 demonstrate evidence for positive selection on at least one branch of the phylogeny constructed using the novel genome assemblies, with gene ontology terms such as calcium ion signaling, histone binding, and protein acetylation identified as being biased in the set of selected genes.

## METHODS

### Species Identification

Wild *Anopheles* larvae belonging to the species *An. crucians* and *freeborni* were provided to the Rasgon Vector Genetics Laboratory by collaborators at the Brunswick County Vector Control Agency (North Carolina, *An. crucians*) and the Benton County Mosquito Control Agency (Washington state, *An. freeborni*). Larvae were shipped from their collection localities in Whirl-Pak bags (Whirl-Pak, Madison, WI, USA) in water from their collection source, at which point they were placed into pans and reared in the Millennium Sciences Complex Insectary (The Pennsylvania State University, University Park, PA, USA) at 27°C ± 1°C, 12 hour light 12 hour dark diurnal cycle at 80% relative humidity. Ground fish flakes (TetraMin, Melle, Germany) were used to feed larvae, and upon emergence adult mosquitoes were maintained with a 10% sucrose solution. *An. quadrimaculatus* and *albimanus* were taken from colony populations maintained in the Millennium Sciences Complex Insectary initially provided by BEI Resources (Manassas, VA, USA).

Fourth instar larvae were isolated and identified using morphological features under light microscopy according to reference material for *Anopheles* larval identification in “Mosquitoes of Pennsylvania”, and species ranges of identifications were confirmed using ranges published in “Mosquitoes of New York” [11, 12]. Larvae were reared separately after identification according to date of emergence and species. Following emergence species identification was then confirmed in adults using morphological characteristics for adults as described in “Mosquitos of Pennsylvania” [12]. Samples NC crucians 5 – 9 were originally identified as *An. punctipennis* by morphology, but mitochondrial assemblies placed them as *An. crucians* (see Results).

### Genome Assembly

DNA was extracted from individual mosquitos using the Qiagen DNEasy Blood and Tissue Kit according to the manufacturer’s suggested protocols. Extracted DNA underwent library preparation and genome sequencing at the New York University Langone Genome Technology Center (New York, NY, USA) using a NovaSeq 6000 with 150 bp paired end sequencing to ∼30X coverage per sample. Raw sequencing reads had adapters trimmed and low-quality bases removed using Trimmomatic read trimming software with base settings [13]. Velvet was run on trimmed and paired reads for each sample with k-mer length of 85 bp, minimum coverage of 5x and expected coverage of 30x to generate the novel genome assemblies [14]. Genome completeness was measured using BUSCO with the insecta_odb10 database to scan for complete conserved single-copy orthologs present within the novel assemblies [15].

Mitochondrial genomes were assembled from interleaved raw sequencing reads for each sample using the mitoBIM pipeline [16] and were then aligned using BLAST against a database of cytochrome c oxidase subunit 1 sequences collected from NCBI to determine closest species assignment for the mitochondrial sequences (Supplementary Table 1) [17]. Augustus de-novo gene prediction software was used to identify potential proteins within the novel assemblies using a training set of 5876 full length coding sequences from the *An. gambiae* PEST strain AgamP3 (GCA_000005575) genome assembly and functional prediction on the potential gene sets was achieved with InterProScan [18, 19].

### Phylogenetics and Scanning for Signatures of Selection

To identify single copy-orthologs for use in creating a phylogeny for the novel genomes and for identifying signatures of selection, Exonerate protein2genome was run using proteins present in the *An. gambiae* PEST strain UP000007062 proteome published on uniProt against novel genome assemblies [20, 21]. Exonerate hits that had scores above 1000, were represented at least 80% of full length in the novel genomes, were not duplicated in any genome, and were present in all genomes considered were kept for further analysis (Supplementary File 1, Supplementary Table 2). Multiple sequence alignment (MSA) for all orthologs were generated using MUSCLE and pal2nal was used to convert the MSA for each ortholog into an in-frame codon alignment [22, 23]. Concatenated in-frame codon alignments for all orthologs were used to create a consensus phylogenetic tree using iqTree, then individual gene-trees were made for each ortholog and compared with the consensus tree to determine gene concordance and site concordance factors for each node of the consensus tree [24]. HyPhy software was used to run Mixed Effects Model of Evolution (MEME) to identify signatures of episodic positive selection occurring along any branch of the consensus phylogeny for all orthologs tested, and adaptive Branch-Site Random Effects Likelihood (aBSREL) to identify if positive selection occurred along any specific proportion of branches for the orthologs in question [25, 26].

## RESULTS/DISCUSSION

### Genome Assembly

We created novel genome assemblies from wild-collected *An. crucians* and *An. freeborni* and from colony populations of *An. quadrimaculatus* and *An. albimanus*. For the natural samples representing *An. crucians*, we collected whole genome sequences from 6 individuals (NC crucians 2, 3, 4, 5, 6, and 8), and two *An. freeborni* were sequenced (WA freeborni 1 and 2) (Table 1). Of the colony *An. quadrimaculatus* and *An. albimanus*, 2 individuals each were sent for whole genome sequencing (COL albi 1 and 2, and COL quad 3 and 4).

**Table 1:** Information related to natural mosquito collections.

| State | Molecular Species ID | Specimen Number | ID | Date Received | DPE | DNA Concentration (ng/uL - 100 uL Total Volume) | Collection GPS | Proportion of mosquito for DNA extraction |
| --- | --- | --- | --- | --- | --- | --- | --- | --- |
| NC | *An. crucians* | 2 | NC crucians 2 | 3/April/2019 | 6 | 2.66 | 34.1697, -78.1688 | 1/10 mosquito |
| NC | *An. crucians* | 3 | NC crucians 3 | 3/April/2019 | 6 | 4.15 | 34.1697, -78.1688 | 1/10 mosquito |
| NC | *An. crucians* | 4 | NC crucians 4 | 3/April/2019 | 9 | 3.87 | 34.1697, -78.1688 | 1/10 mosquito |
| NC | *An. crucians* | 5 | NC crucians 5 | 3/April/2019 | 3 | 3.63 | 34.1697, -78.1688 | 1/10 mosquito |
| NC | *An. crucians* | 6 | NC crucians 6 | 3/April/2019 | 4 | 2.21 | 34.1697, -78.1688 | 1/10 mosquito |
| NC | *An. crucians* | 8 | NC crucians 8 | 3/April/2019 | 6 | 4.32 | 34.1697, -78.1688 | 1/10 mosquito |
| WA | *An. freeborni* | 1 | WA freeborni 1 | 23/May/2019 | 5 | 8.33 | 46.339903, -119.379405 | half mosquito |
| WA | *An. freeborni* | 2 | WA freeborni 2 | 23/May/2019 | 5 | 6.1 | 46.339903, -119.379405 | half mosquito |
| COLONY | *An. quadrimaculatus* | 3 | COL quad 3 |  |  | 40.15 |  | whole mosquito |
| COLONY | *An. quadrimaculatus* | 4 | COL quad 4 |  |  | 42.78 |  | whole mosquito |
| COLONY | *An. albimanus* | 1 | COL albi 1 |  |  | 17.41 |  | whole mosquito |
| COLONY | *An. albimanus* | 2 | COL albi 2 |  |  | 14.46 |  | whole mosquito |

Table 2 displays genome sizes, N50, and BUSCO scores for all samples sequenced. *An. crucians* had the largest genome size of those sequenced in this study with 310.8 Mb average size per assembly, *An. freeborni* had an average genome size of 270.05 Mb, *An. quadrimaculatus* had an average genome size of 229.95 Mb, and *An. albimanus* had the smallest genome size of those sampled with an average assembly size of 170.85 Mb. N50 was highest for colony sourced samples which is likely due to lower genetic diversity within colony populations vs natural populations, and ranged from 213.5 Kb for *An. albimanus* sample COL albi 1, to 25.9 Kb for *An. crucians* sample NC crucians 5 (Table 2). *An. albimanus* is the only species represented in this study that already has a genome assembly published, and they report a genome size of 170.85 Mb [27], very similar to our results. BUSCO scores suggest all assemblies are relatively complete with between 98.3% and 99.5% single-copy ortholog representation from the BUSCO insecta_odb10 dataset [15].

**Table 2:** Genome sizes, N50, and BUSCO scores for all samples sequenced.

| Sample | Species | Assembly Size (MB) | N50 (KB) | Single Copy BUSCO | Duplicated BUSCO | Fragmented BUSCO | Missing BUSCO | Total BUSCO |
| --- | --- | --- | --- | --- | --- | --- | --- | --- |
| COL albi 1 | *An. albimanus* | 170.3 | 213.5 | 1346 | 13 | 2 | 6 | 1367 |
| COL albi 2 | *An. albimanus* | 171.4 | 200.1 | 1348 | 10 | 2 | 7 | 1367 |
| COL quad 3 | *An. quadrimaculatus* | 229.2 | 110.3 | 1322 | 32 | 5 | 8 | 1367 |
| COL quad 4 | *An. quadrimaculatus* | 230.7 | 100.1 | 1316 | 28 | 10 | 13 | 1367 |
| NC crucians 2 | *An. crucians* | 309.4 | 30.0 | 1315 | 29 | 10 | 13 | 1367 |
| NC crucians 3 | *An. crucians* | 304.2 | 34.0 | 1323 | 26 | 9 | 9 | 1367 |
| NC crucians 4 | *An. crucians* | 312.8 | 35.9 | 1320 | 25 | 12 | 10 | 1367 |
| NC crucians 5 | *An. crucians* | 321.6 | 25.9 | 1318 | 31 | 11 | 7 | 1367 |
| NC crucians 6 | *An. crucians* | 308.1 | 35.6 | 1320 | 25 | 8 | 14 | 1367 |
| NC crucians 8 | *An. crucians* | 308.8 | 39.3 | 1319 | 30 | 10 | 8 | 1367 |
| WA freeborni 1 | *An. freeborni* | 268.7 | 53.6 | 1335 | 12 | 3 | 17 | 1367 |
| WA freeborni 2 | *An. freeborni* | 274.1 | 57.0 | 1339 | 14 | 4 | 10 | 1367 |

A trend for high numbers of predicted genes was observed in the natural genomes, with as many as 40 thousand predicted for some *An. crucians* samples, and functional prediction using InterProScan determined many of these gene predictions to be classified as intrinsically disordered proteins [19]. Many of these predicted genes were identified as intrinsically disordered proteins in the novel assemblies occurred as the beginning or end of a contig, and with the shorter contig lengths in natural vs colony sourced genomes. It is likely that the large number of poorly categorized proteins in the natural assemblies represent full length proteins which have been fragmented by the lower quality of the assembly. Future long read resequencing of these species will help to address this issue.

### Phylogenetics

A total of 790 single copy orthologs from the *An. gambiae* PEST strain proteome on UniProt were identified that were shared between all novel genome assemblies (Supplementary File 1, Supplementary Table 1) [20]. A maximum likelihood phylogeny was created using a concatenated alignment for a selection of 673 orthologs that were shared with *Aedes aegypti* which acted as an outgroup. Individual trees were made for each ortholog for calculation of site and gene concordance factors using the concatenated maximum likelihood phylogeny as a reference (Figure 1).

**Figure 1:**
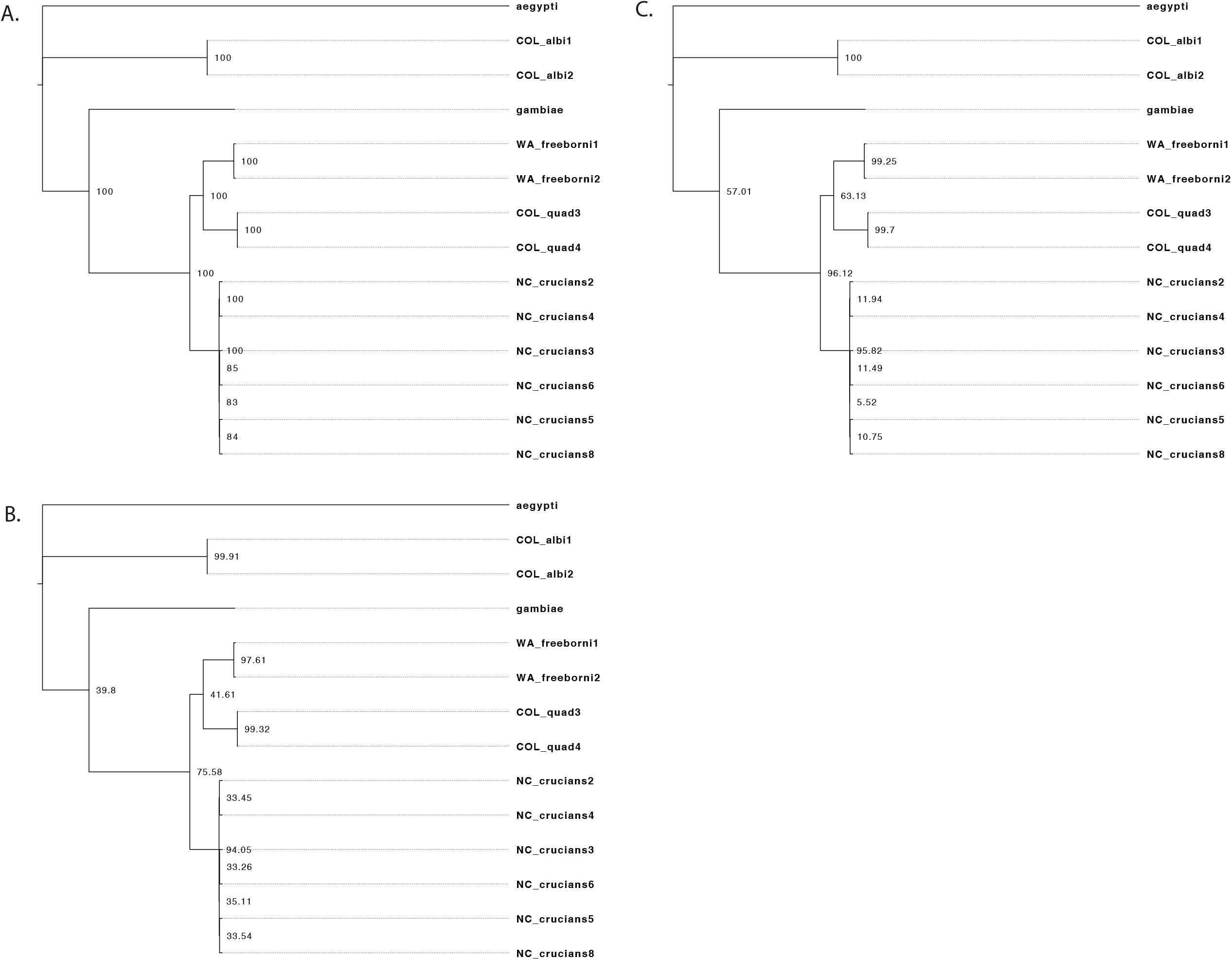
Maximum liklihood phylogenies with bootstrap values (A.), gene concordance factors (B.), and site concordance factors (C.) displayed on corresponding nodes.

*An. albimanus* and *An. gambiae* are placed on their own independent branches, while *An. freeborni, An. quadrimaculatus*, and *An. crucians* are on a branch together with 100% bootstrap support. *An. freeborni* and *An. quadrimaculatus* diverged with 100% bootstrap support into independent branches, while *An. crucians* diverges into its own branch with 100% bootstrap support, however the topology within this branch was not well supported. There are nodes within the branch that makes the *An. crucians* portion of the phylogeny, each containing 2 samples with 2 gene concordance factors of 11.94, 11.49, and 10.75.

Mitochondrial genomes were assembled from the raw sequencing data collected for each sample, and these genomes were compared to a database of cytochrome c oxidase subunit 1 sequences representing *An. gambiae, quadrimaculatus, albimanus, freeborni, crucians*, and *punctipennis* with BLAST to determine species identification (Supplementary Table 1) [17]. The potential for admixture was considered between *An. crucians* and *punctipennis* as this has been observed previously for *An. crucians* and *bradleyi* [28], but analysis of genome variants with ADMIXTURE and did not identify any evidence for hybridization between samples from the samples which could have represented the two species or confidently place them into two groups along species lines (Figure 2) [29]. PCA on genomic variants identified within the ortholog set demonstrates that the *An. crucians* branch separates into two loose groups, but they cluster much closer together compared with the other species represented within this study (Figure 3). The evidence suggests that instead the branch for *An. crucians* represents a single species, not 2 species or a combination of 2 species and hybridized individuals.

**Figure 2:**
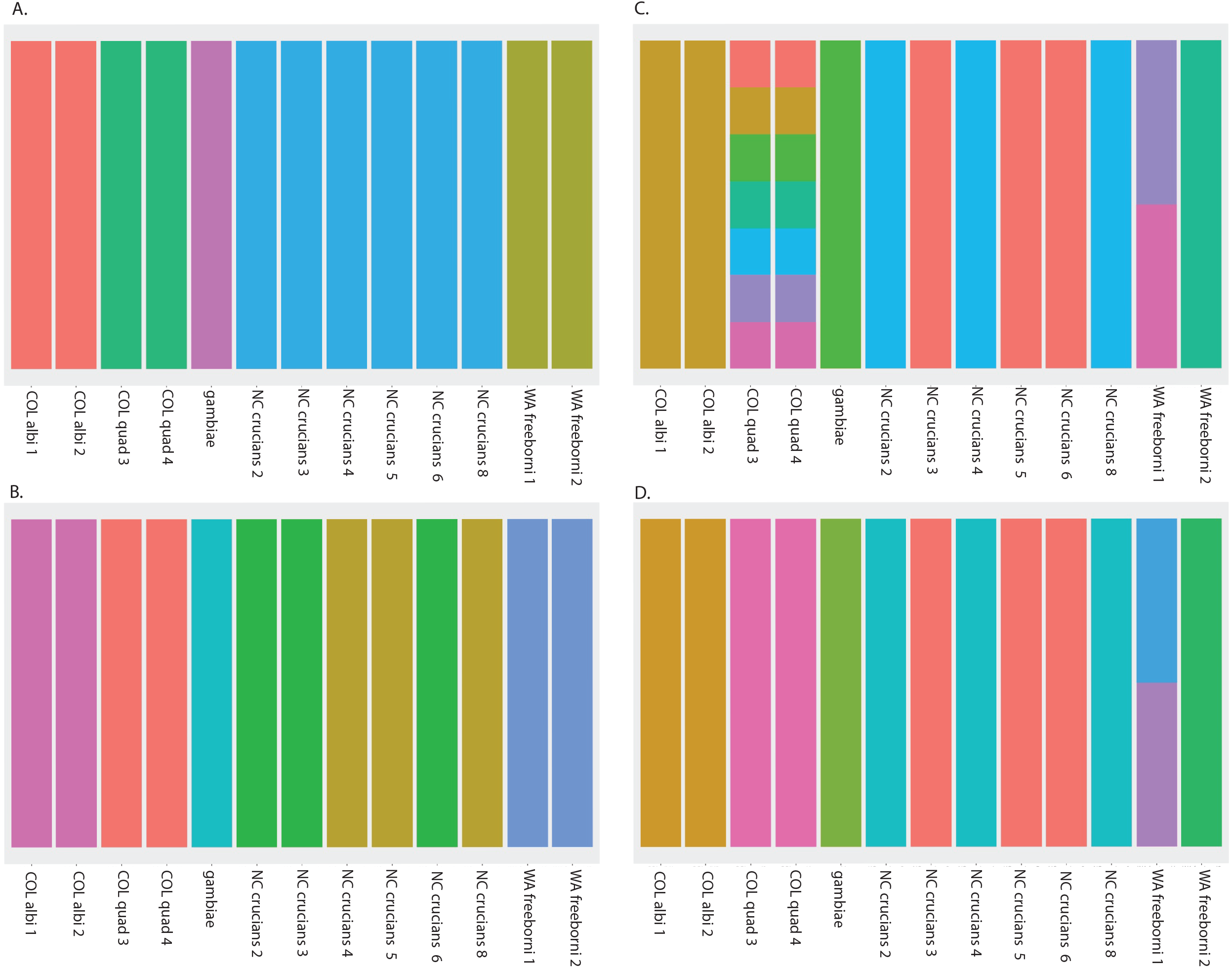
Admixture plots created using ADMIXTURE software with inferred number of populations (K) equal to 5 (A), 6 (B) 7 (C), and 8 (D). Y-axis shows the proportion of a sample’s genome from 0% to100%, X axis has the novel genome assemblies investigated in this study and *An. gambiae* as an outgroup, and the color of the bar for each samle represents the inferred populations representing that sample’s genome. The color’’s height on the Y axis represents the proportion of ancestry that population represents in the corresponding genome assembly.

**Figure 3:**
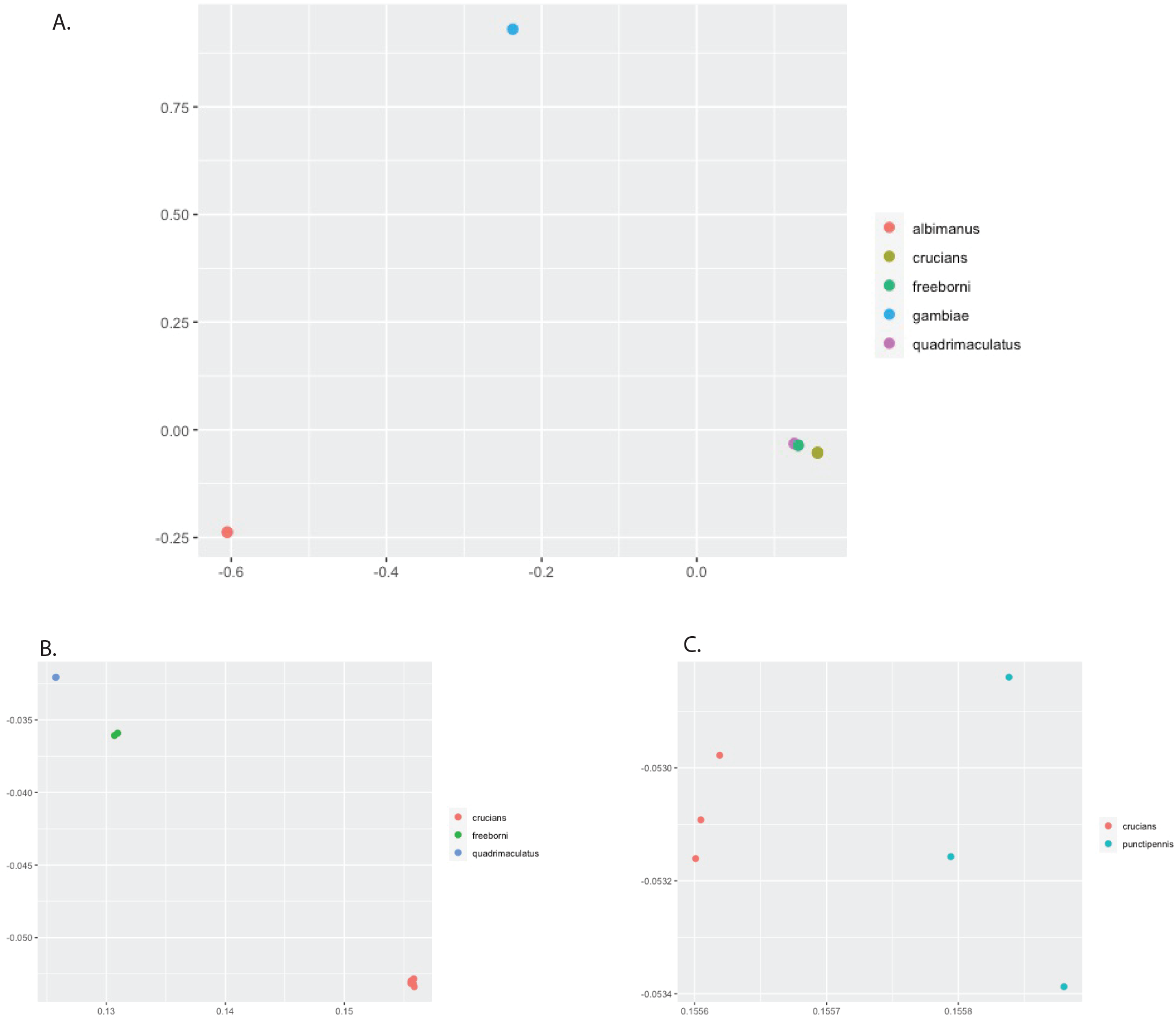
Principal components analysis (PCA) of variants in the 790 identified single copy orthologs shared between all novel assemblies and the *An. gambiae* PEST genome assembly and color coded by species. A shows all samples, B isolates the more closely related group of *An. quadrimaculatus, An. crucians*, and *An. freeborni*, while C isolates *An. crucians* and deliniates between the morphological IDs for *An. crucians* and *An. punctipennis*.

### Evidence for Selection

dN/dS based analyses were performed using HyPhy to analyze signatures of selection in alignments of the full set of 790 single copy orthologs identified in all newly sequenced samples with a reduced tree not containing the *Aedes aegypti* outgroup (Supplementary Figure 1). Signatures of episodic positive selection at individual sites in the orthologs were identified using MEME [25], and selection of orthologs along a proportion of branches within the phylogeny using aBSREL [26]. Of the 790 orthologs considered, MEME identified at least one site under episodic positive selection in 525 orthologs in at least one branch of the phylogeny (Supplementary table 3). Gene ontology analysis using g:Profiler on the genes under episodic selection identified molecular functions related to histone binding and protein acetylation, and cellular components of the cytoplasm and endomembrane complex (Supplementary Table 4) [30]. aBSREL identifies genes under selection in any branch within the phylogeny, and a more diverse set of terms is identified 289 unique orthologs regulated in at least 1 of the 19 nodes identified in the phylogeny (Supplementary Table 3). Gene ontology analysis using g:Profiler on terms regulated in any particular branch identified genes involved with water ion transport and glutamic acid ligation selected for nodes leading to *An. quadrimaculatus* samples (Supplementary Table 4) [30]. Nodes leading to all samples but *An. gambiae* are biased for terms related to cellular components of DNA driven RNA polymerase complex, peroxisomes, intracellular transferase complex, and protein complexes. NC *An. crucians* sample 3 had biased biological process terms related to calcium ion transport, as well as broader terms such as RNA processing and capping, gene expression, and cyclic compound metabolic processing.

## Conclusion

Limiting the spread of vector borne disease can be improved by a deep understanding of the biology of the vector species of interest, and increasing our understanding of vector genetics is essential for progress in eliminating vector borne pathogens. Phylogenetics of novel genome assemblies for *An. freeborni, crucians, albimanus*, and *quadrimaculatus* show that *An. crucians* groups closely with *An. quadrimaculatus* and *freeborni* separate from *An. albimanus*, the only South American species represented, which forms its own clade. Scanning for signatures of selection shows a diverse set of genes being selected for along different branches of the phylogeny, most notably calcium ion transport in *An. crucians* and *quadrimaculatus*. These novel genome sequences will be useful in developing our understanding of the diverse biological traits that drive vectorial capacity in anophelines.

## Supporting information

Supplementary Othologous sequence data fasta file

## DATA AVAILABILITY

Genome sequences are available through NCBI under Bioproject ID PRJNA758127, which encompasses reference numbers SAMN21013496 - SAMN21013507. File S1 contains fasta sequences for the 790 orthologs used in the phylogenetic and hyPhy analyses. Table S1 contains BLAST results against the mitochondrial genome assemblies. Table S2 contains the UniProt and NCBI IDs corresponding to the orthologs used in the phylogenetic and hyPhy analyses. Table S3 contains detailed output for hyPhy analysis. Table S4 contains detailed results for gProfiler analysis on genes identified as being selected for in hyPhy analysis. Figure S1 demonstrates a phylogeny created using iqTree without *Aedes aegypti* as an outgroup, used for running hyPhy analysis.

## ACKNOWLEDGEMENTS

This work was supported by NIH grants R01AI128201, R01AI116636, NSF BIO grant 1645331, USDA Hatch funds (Project #PEN04769), and a grant with the Pennsylvania Department of Health using Tobacco Settlement Funds to JLR. CAH was supported by an NSF graduate research fellowship program award 2018258101.

## Figure legends

Figure 1: Maximum likelihood phylogenies with bootstrap values (A.), gene concordance factors (B.), and site concordance factors (C.) displayed on corresponding nodes.

Figure 2: Admixture plots created using ADMIXTURE software with inferred number of populations (K) equal to 5 (A), 6 (B), 7 (C), and 8 (D). Y-axis shows the proportion of a sample’s genome from 0% to100%, X axis has the novel genome assemblies investigated in this study and *An. gambiae* as an outgroup, and the color of the bar for each sample represents the inferred populations representing that sample’s genome. The color ‘s height on the Y axis represents the proportion of ancestry that population represents in the corresponding genome assembly.

Figure 3: Principal components analysis (PCA) of variants in the 790 identified single copy orthologs shared between all novel assemblies and the *An. gambiae* PEST genome assembly and color coded by species. A shows all samples, B isolates the more closely related group of *An. quadrimaculatus, An. crucians*, and *An. freeborni*, while C isolates and delineates between the morphological IDs for *An. crucians* and *An. punctipennis*.

## REFERENCES

1. Franz A, Kantor A, Passarelli A, Clem R. Tissue Barriers to Arbovirus Infection in Mosquitoes. Viruses. 2015;7(7):3741–3767. doi:10.3390/v7072795

2. Vector-borne diseases. https://www.who.int/news-room/fact-sheets/detail/vector-borne-diseases. Accessed March 2, 2021.

3. Sinka ME, Rubio-Palis Y, Manguin S, et al. The dominant Anopheles vectors of human malaria in the Americas: occurrence data, distribution maps and bionomic précis. Parasit Vectors. 2010;3(1):72. doi:10.1186/1756-3305-3-72

4. Rezza G, Chen R, Weaver SC. O’nyong-nyong fever: a neglected mosquito-borne viral disease. Pathog Glob Health. 2017;111(6):271–275. doi:10.1080/20477724.2017.1355431

5. Yadav P, Barde P, Singh D, Mishra A, Mourya D. EXPERIMENTAL TRANSMISSION OF CHIKUNGUNYA VIRUS BY ANOPHELES STEPHENSI MOSQUITOES. Acta Virol. 2003;47:45–47. http://www.elis.sk/download_file.php?product_id=8&session_id=jc2533t2t4ld65j13rlddb4r55. Accessed February 24, 2019.

6. Brustolin M, Pujhari S, Henderson CA, Rasgon JL. Anopheles mosquitoes may drive invasion and transmission of Mayaro virus across geographically diverse regions. Christofferson RC, ed. PLoS Negl Trop Dis. 2018;12(11):e0006895. doi:10.1371/journal.pntd.0006895

7. Chakraborty M, Ramaiah A, Adolfi A, et al. Hidden genomic features of an invasive malaria vector, Anopheles stephensi, revealed by a chromosome-level genome assembly. BMC Biol. 2021;19(1):1–16. doi:10.1186/s12915-021-00963-z

8. Miles A, Harding NJ, Bottà G, et al. Genetic diversity of the African malaria vector anopheles gambiae. Nature. 2017;552:96–100. doi:10.1038/nature24995

9. Holt RA, Mani Subramanian G, Halpern A, et al. The genome sequence of the malaria mosquito Anopheles gambiae. Science (80-). 2002;298(5591):129–149. doi:10.1126/science.1076181

10. Pritchard AE, Seabrook EL, Provost MW. The possible endemicity of Anopheles albimanus in Florida. Mosq News. 1946 Dec;6(4):183. PMID: 20292848.

11. Means RG. Mosquitoes of New York: Genera of Culicidae Other than Aedes Occurring in … - Robert G. Means - Google Books. New York: University of the State of New York; 1987. https://books.google.com/books/about/Mosquitoes_of_New_York_Genera_of_Culicid.html?id=yvUhAQAAMAAJ. Accessed March 2, 2021.

12. Darsie RF, Hutchinson ML. THE MOSQUITOES OF PENNSYLVANIA Identification of Adult Females and Fourth Instar Larvae, Geographical Distribution, Biology and Public Health Importance.; 2009.

13. Bolger AM, Lohse M, Usadel B. Trimmomatic: a flexible trimmer for Illumina sequence data. Bioinformatics. 2014;30(15):2114–2120. doi:10.1093/bioinformatics/btu170

14. Zerbino DR. Using the Velvet de novo assembler for short-read sequencing technologies. Curr Protoc Bioinforma. 2010;CHAPTER(SUPPL. 31):Unit. doi:10.1002/0471250953.bi1105s31

15. Simão FA, Waterhouse RM, Ioannidis P, Kriventseva E V., Zdobnov EM. BUSCO: assessing genome assembly and annotation completeness with single-copy orthologs. Bioinformatics. 2015;31(19):3210–3212. doi:10.1093/bioinformatics/btv351

16. Hahn C, Bachmann L, Chevreux B. Reconstructing mitochondrial genomes directly from genomic next-generation sequencing reads - A baiting and iterative mapping approach. Nucleic Acids Res. 2013;41(13):e129–e129. doi:10.1093/nar/gkt371

17. Altschul SF, Gish W, Miller W, Myers EW, Lipman DJ. Basic local alignment search tool. J Mol Biol. 1990;215(3):403–410. doi:10.1016/S0022-2836(05)80360-2

18. Stanke M, Diekhans M, Baertsch R, Haussler D. Using native and syntenically mapped cDNA alignments to improve de novo gene finding. Bioinformatics. 2008;24(5):637–644. doi:10.1093/bioinformatics/btn013

19. Jones P, Binns D, Chang HY, et al. InterProScan 5: Genome-scale protein function classification. Bioinformatics. 2014;30(9):1236–1240. doi:10.1093/bioinformatics/btu031

20. Bateman A, Martin MJ, Orchard S, et al. UniProt: The universal protein knowledgebase in 2021. Nucleic Acids Res. 2021;49(D1):D480–D489. doi:10.1093/nar/gkaa1100

21. Slater GSC, Birney E. Automated generation of heuristics for biological sequence comparison. BMC Bioinformatics. 2005;6(1):31. doi:10.1186/1471-2105-6-31

22. Edgar RC. MUSCLE: Multiple sequence alignment with high accuracy and high throughput. Nucleic Acids Res. 2004;32(5):1792–1797. doi:10.1093/nar/gkh340

23. Suyama M, Torrents D, Bork P. PAL2NAL: robust conversion of protein sequence alignments into the corresponding codon alignments. doi:10.1093/nar/gkl315

24. Minh BQ, Schmidt HA, Chernomor O, et al. IQ-TREE 2: New Models and Efficient Methods for Phylogenetic Inference in the Genomic Era. Teeling E, ed. Mol Biol Evol. 2020;37(5):1530–1534. doi:10.1093/molbev/msaa015

25. Murrell B, Wertheim JO, Moola S, Weighill T, Scheffler K, Kosakovsky Pond SL. Detecting individual sites subject to episodic diversifying selection. PLoS Genet. 2012;8(7):1002764. doi:10.1371/journal.pgen.1002764

26. Smith MD, Wertheim JO, Weaver S, Murrell B, Scheffler K, Kosakovsky Pond SL. Less Is More: An Adaptive Branch-Site Random Effects Model for Efficient Detection of Episodic Diversifying Selection. Mol Biol Evol. 2015;32(5):1342–1353. doi:10.1093/molbev/msv022

27. Neafsey DE, Waterhouse RM, Abai MR, et al. Highly evolvable malaria vectors: The genomes of 16 Anopheles mosquitoes. Science (80-). 2015;347(6217). doi:10.1126/science.1258522

28. Kreutzer RD, Kitzmiller JB. Hybridization Between Anopheles crucians and Anopheles bradleyi. Evolution (N Y). 1971;25(1):195. doi:10.2307/2406511

29. Alexander DH, Lange K. Enhancements to the ADMIXTURE algorithm for individual ancestry estimation. BMC Bioinformatics. 2011;12(1):246. doi:10.1186/1471-2105-12-246

30. Uri Reimand J, Arak T, Adler P, et al. g:Profiler-a web server for functional interpretation of gene lists (2016 update). Nucleic Acids Res. 2016;(1). doi:10.1093/nar/gkw199

