## Supplementary Figure 1 for "Novel genome assemblies and Evolutionary Dynamics of North American *Anopheles* mosquitoes"

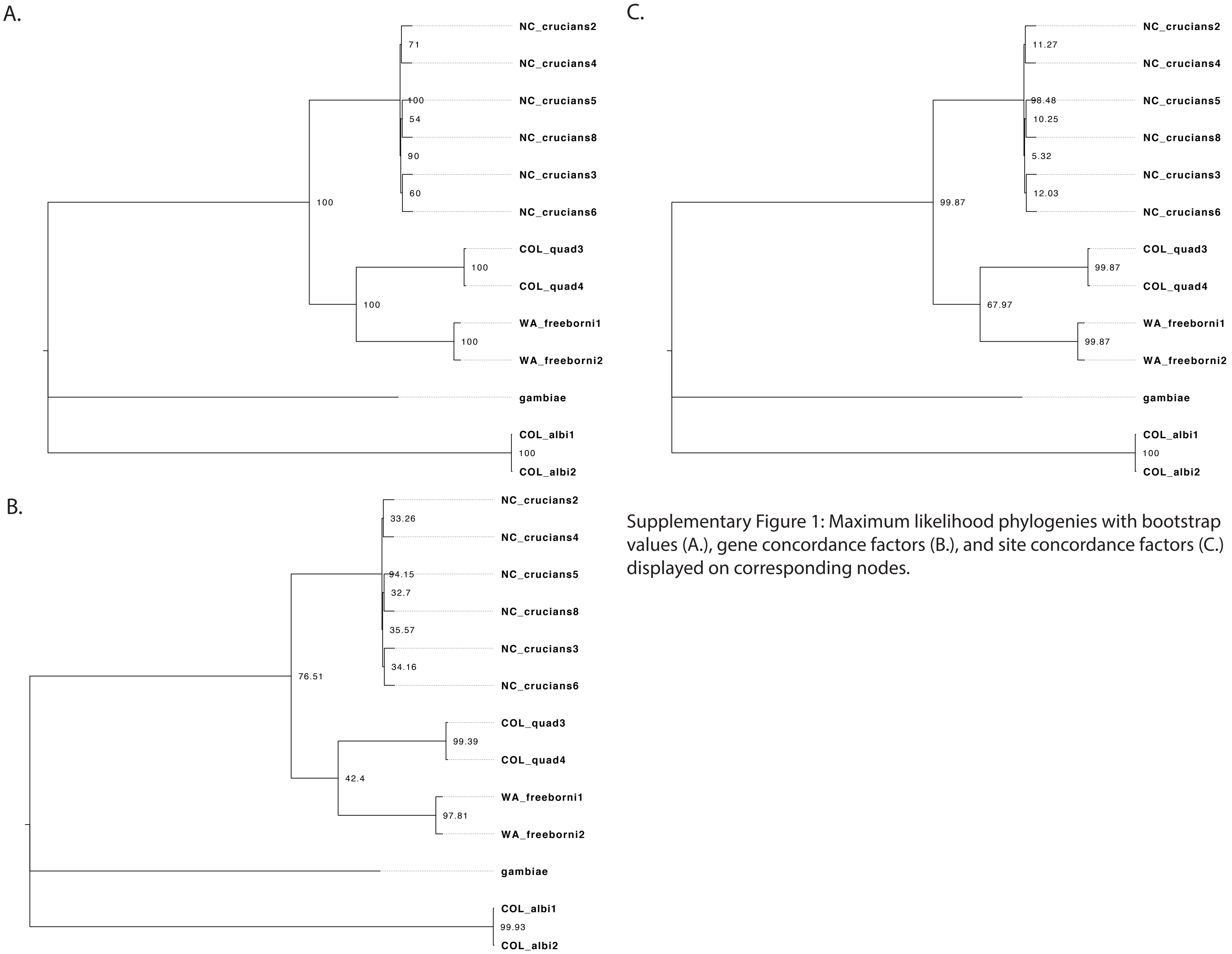


Supplementary Figure 1: Maximum likelihood phylogenies with bootstrap values (A.), gene concordance factors (B.), and site concordance factors (C.) displayed on corresponding nodes.
